# Transcriptional activity differentiates families of Marine Group II *Euryarchaeota* in the coastal ocean

**DOI:** 10.1101/2020.09.16.299958

**Authors:** Julian Damashek, Aimee Oyinlade Okotie-Oyekan, Scott Michael Gifford, Alexey Vorobev, Mary Ann Moran, James Timothy Hollibaugh

**Author notes:** **Corresponding author:** Julian Damashek, Utica College, 1600 Burrstone Road, Utica, NY 13502.

## Abstract

Marine Group II *Euryarchaeota* (*Candidatus* Poseidoniales), abundant but yet-uncultivated members of marine microbial communities, are thought to be (photo)heterotrophs that metabolize dissolved organic matter (DOM) such as lipids and peptides. However, little is known about their transcriptional activity. We mapped reads from a metatranscriptomic time series collected at Sapelo Island (GA, USA) to metagenome-assembled genomes to determine the diversity of transcriptionally-active *Ca*. Poseidoniales. Summer metatranscriptomes had the highest abundance of *Ca*. Poseidoniales transcripts, mostly from the O1 and O3 genera within *Ca*. Thalassarchaeaceae (MGIIb). In contrast, transcripts from fall and winter samples were predominantly from *Ca*. Poseidoniaceae (MGIIa). Genes encoding proteorhodopsin, membrane-bound pyrophosphatase, peptidase/proteases, and part of the β-oxidation pathway were highly transcribed across abundant genera. Highly transcribed genes specific to *Ca*. Thalassarchaeaceae included xanthine/uracil permease and receptors for amino acid transporters. Enrichment of *Ca*. Thalassarchaeaceae transcript reads related to protein/peptide, nucleic acid, and amino acid transport and metabolism, as well as transcript depletion during dark incubations, provided further evidence of heterotrophic metabolism. Quantitative PCR analysis of South Atlantic Bight samples indicated consistently abundant *Ca*. Poseidoniales in nearshore and inshore waters. Together, our data suggest *Ca*. Thalassarchaeaceae are important photoheterotrophs potentially linking DOM and nitrogen cycling in coastal waters.

## INTRODUCTION

Since the initial discovery of Marine Group II (MGII) *Euryarchaeota* [1,2], definitive determination of their physiology and ecological roles has remained challenging due to the lack of a cultivated isolate. Nonetheless, as data describing MGII distributions throughout the oceans have increased, several patterns have emerged: MGII are often highly abundant in the euphotic zone and in coastal waters, can reach high abundance following phytoplankton blooms, and are largely comprised of two subclades, MGIIa and MGIIb [3,4]. Early metagenomic studies provided the first evidence that MGII may be aerobic (photo)heterotrophs [5–7], a hypothesis supported by incubation experiments [8–10] and by the gene content of diverse metagenome-assembled genomes (MAGs) [11–15]. Two recent studies deepened our understanding of the phylogenomics and metabolic potential of MGII by analyzing hundreds of MAGs, highlighting clade-specific differences in genomic potential for transport and degradation of organic molecules, light harvesting proteorhodopsins, and motility [16,17]. Here, we refer to MGII as the putative order *“Candidatus* Poseidoniales,” MGIIa and MGIIb as the putative families “*Ca*. Poseidoniaceae” and “*Ca*. Thalassarchaeaceae,” respectively, and putative genera as specified by Rinke et al. [16]. We occasionally use “MGIIa” and “MGIIb” for consistency with previous literature.

Metatranscriptomics is one strategy for gleaning information about microbial activity in the environment. *Ca*. Poseidoniales transcripts can be abundant in marine metatranscriptomes, suggesting transiently high transcriptional activity [18,19]. When metatranscriptome reads from the Gulf of Aqaba were mapped to metagenomic contigs from the Mediterranean Sea, genes involved in amino acid transport, carbon metabolism, and cofactor synthesis were highly transcribed in the aggregate euryarchaeal community [20,21]. In another study, mapping deep-sea metatranscriptome reads to novel *Ca*. Poseidoniales MAGs indicated transcription of genes related to protein, fatty acid, and carbohydrate transport and metabolism, likely fueling aerobic heterotrophy [15]. Finally, a metaproteomics study found abundant euryarchaeal transport proteins for L-amino acids, branched-chain amino acids, and peptides throughout the Atlantic Ocean [22]. Despite these advances, little is known about similarities or differences in gene transcription between *Ca*. Poseidoniales and Thalassarchaeaceae.

We report MAG-resolved metatranscriptomic analyses of *Ca*. Poseidoniales in coastal waters near Sapelo Island (GA, USA). Prior work suggested *Ca*. Poseidoniales are sporadically active at Sapelo Island [23] and may comprise the majority of archaea in mid-shelf surface waters of the South Atlantic Bight (SAB) [24]. Since other studies thoroughly described the genomic content of *Ca*. Poseidoniales MAGs, our focus instead was determining which clades were transcriptionally active and identifying highly or differentially transcribed genes. We used two Sapelo Island MAGs [25] combined with recent *Ca*. Poseidoniales MAG collections [16,17] to competitively recruit reads from a metatranscriptomic time series [26] and an incubation experiment [23] to determine which clades were active over time. We then used representative MAGs from highly active genera to determine which *Ca*. Poseidoniales genes were transcribed. Finally, we used quantitative PCR (qPCR) to measure the abundance of *Ca*. Poseidoniales 16S rRNA genes in DNA samples throughout the SAB to assess the prevalence *Ca*. Poseidoniales in this region.

## MATERIALS AND METHODS

### PHYLOGENOMICS

Phylogenomic analyses compared SIMO Bins 19-2 and 31-1 (ref. 25) to previously-reported *Ca*. Poseidoniales MAGs, including those binned from TARA Oceans, Mediterranean Sea, Red Sea, Gulf of Mexico, Guaymas basin, and Puget Sound metagenomes [11,13,15–17,27–32] (Table S1). Average nucleotide identity (ANI) was calculated using fastANI [33] to compare non-redundant MAGs from Tully [17] to 15 Port Hacking MAGs [16] and the two SIMO MAGs; any with ANI <98.5% were added to the non-redundant set. Phylogenomic analysis used sixteen ribosomal proteins [34] within anvi’o v4 (ref. 35). All genomes were converted to contig databases and proteins were identified using HMMER [36], concatenated, aligned using MUSCLE [37], and used to build a phylogenomic tree using FastTree [38] within anvi’o.

### COMPETITIVE READ MAPPING

We used competitive read mapping to determine which *Ca*. Poseidoniales genera were transcriptionally active in Sapelo Island metatranscriptomes. Surface water for all metatranscriptomes was collected seasonally in 2008, 2009, and 2014 from the dock at Marsh Landing, Sapelo Island (31°25′4.08 N, 81°17′43.26 W) as previously described [23,26]. Briefly, 3-8 L of surface water was pumped through a 3-μm pore size prefilter followed by a 0.2-μm pore size collection filter, which was frozen on liquid nitrogen. RNA from the free was extracted from the collection filter as previously described [23,26], including the addition of internal RNA standards to calculate volumetric transcript abundances [39]. Analyses of “field” communities included Gifford et al. metatranscriptomes (iMicrobe Accession <u>CAM_P_0000917</u>) [26] and the T0 metatranscriptomes from Vorobev et al. [23], while “dark incubation” analyses included only Vorobev et al. samples (T0, which were processed immediately after collection, and T24, which were processed following 24 hours of *in situ* incubation in dark containers; NCBI BioProject <u>PRJNA419903</u>). Temperature, salinity, dissolved oxygen, pH, and turbidity data corresponding to metatranscriptome sampling times were downloaded from the NOAA National Estuarine Research Reserve System website (http://cdmo.baruch.sc.edu; last accessed 16 July 2020).

Contigs from all MAGs used for phylogenomics (Table S1) were used as a read mapping database using Bowtie2 v.2.2.9 (ref. 40) with the “very-sensitive” flag. Samtools v.1.3.1 (ref. 41) was used to index BAM files, which were profiled and summarized in anvi’o. Contig genus identity was imported to the anvi’o contig database as an external collection. The number of transcripts L^−1^ was calculated by scaling the number of mapped reads by the volume of water filtered and the recovery of internal standards (reported in [23,26]) as previously described [39]. Seasonal transcript abundances were compared using a one-way ANOVA test in R [42] with data log-transformed as necessary to improve normality. When ANOVA results were significant, groupings were defined post-hoc with Tukey’s Honest Significant Difference (HSD) test using the agricolae R package [43].

Non-metric multidimensional scaling (NMDS) analysis of metatranscriptome hits was conducted using the vegan R package [44]. NMDS input was a distance matrix constructed by Hellinger-transforming the table of transcript hits and calculating Euclidean distance between samples [45]. Genus vectors were calculated using the *envfit* command. Significance of groupings were tested by permutational multivariate analysis of variance (*adonis* command) with 999 permutations.

### MAG-SPECIFIC ANNOTATION AND TRANSCRIPT ANALYSIS

Gene-specific analyses focused on three MAGs: two from the SIMO collection (SIMO Bin 19-2, Genbank: <u>VMDE00000000</u>; SIMO Bin 31-1, <u>VMBU00000000</u>; [25]) and one previously binned from Red Sea metagenomes (RS440, <u>PBUZ00000000</u>; [27]). These MAGs represented genera O1, O3, and M, respectively, which were highly abundant in metatranscriptomes (see Fig. 1). RS440 was selected due to a high number of transcripts recruited when genus M was abundant (data not shown).

**Fig. 1.**
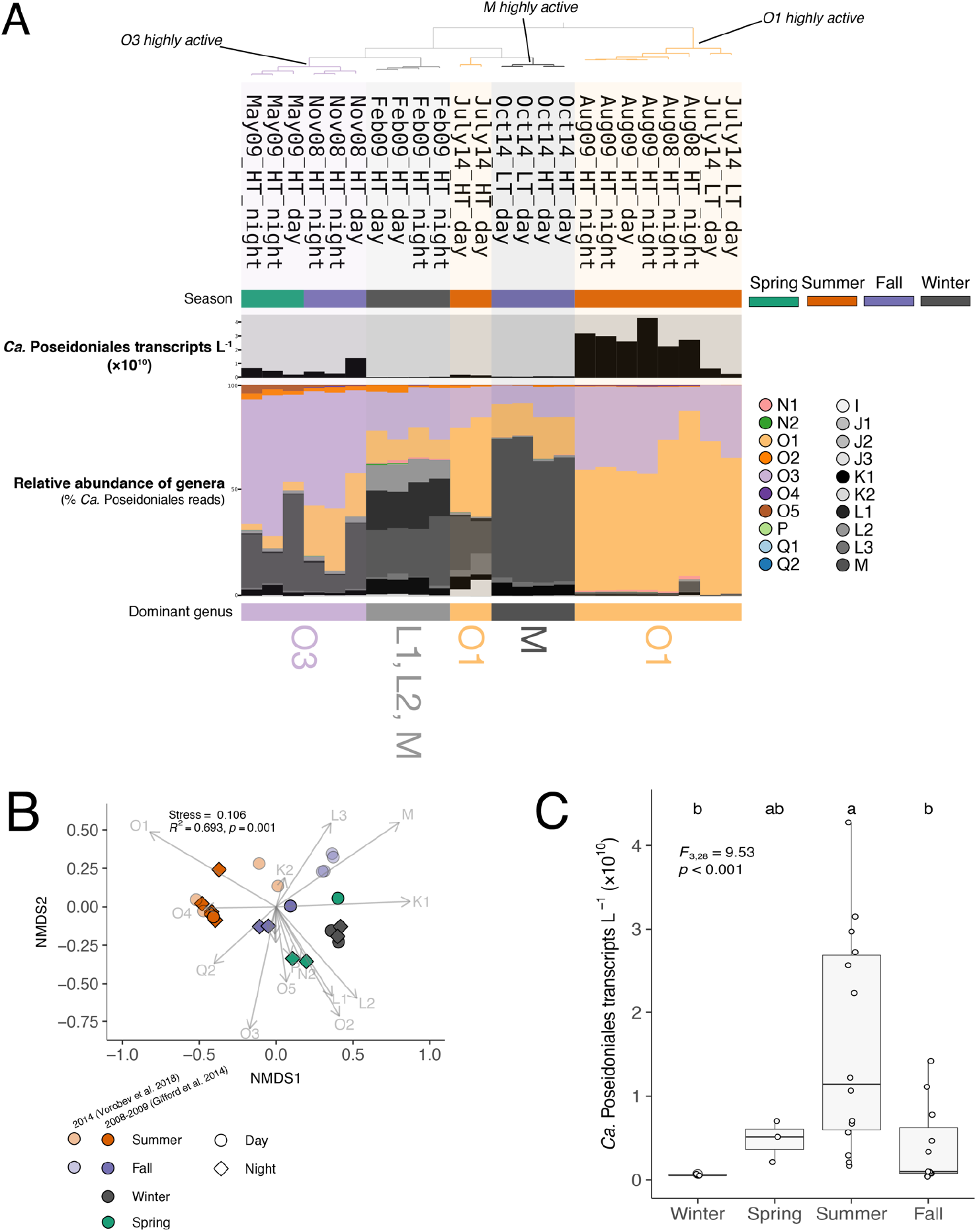
**A)** Relative abundance of *Ca*. Poseidoniales genera in Sapelo Island metatranscriptomes (*n*=24). The dendrogram (top) shows grouping by similarity. Season is indicated by color beneath sample names. The bar chart shows the abundance of *Ca*. Poseidoniales transcripts L^−1^ and the stacked bar charts show the relative abundance of genera (% total *Ca*. Poseidoniales transcripts), colored by genus. Dominant genera are indicated below the stacked bar chart. “Highly active” samples for each genus are marked and were used for analysis of differential transcription. **B)** NMDS of *Ca*. Poseidoniales metatranscriptome reads using a Hellinger distance matrix of genera relative abundances. Each point represents a sample, with color and shape denoting season and day/night, respectively. Shading indicates sampling year and original study. Arrows represent vectors for each genus. NMDS stress and the results of a PERMANOVA analyzing sums of squares by season are indicated. **C)** Boxplots of *Ca*. Poseidoniales reads L^−1^, grouped by season (winter, *n*=4; spring, *n*=3; summer, *n*=10; fall, *n*=7). Values from individual metatranscriptomes are overlain. Results of an ANOVA are indicated; letters at the top indicate post-hoc groups according to Tukey’s HSD test.

MAGs were annotated using the archaeal database in Prokka v.1.13 (ref. 46), using DIAMOND [47] to search against all orthologous groups in eggNOG-mapper v.1 (refs. 45,46), and using the BlastKOALA portal (https://www.kegg.jp/blastkoala/, last accessed 6 March 2019) [48]. Putative genes for carbohydrate-active enzymes, peptidases, and membrane transport proteins were identified using HMMER searches of dbCAN2 (HMMdb v.7) [49,50], MEROPS v.12.0 (ref. 51), and the Transporter Classification Database [52], respectively.

Transcript reads were mapped to MAGs (combined into a single database such that each read mapped to only one MAG) to identify *Ca*. Poseidoniales genes that were highly or differentially transcribed. Coverage was calculated by profiling BAM files in anvi’o and normalized to coverage per million reads (CPM) by dividing by the total number of reads per sample. For each MAG, “highly transcribed” genes were the 5% of putative genes with the highest median CPM across metatranscriptomes (SIMO Bin 19-2: 63 genes, SIMO Bin 31-1: 70 genes, RS440: 77 genes).

DESeq2 v.3.11 (ref. 53) was used to identify genes from each MAG that were differentially transcribed when each genus was highly transcriptionally active. For each MAG, “treatment” samples in DESeq2 were those where the respective genus recruited ≥50% of *Ca*. Poseidoniales reads from the metatranscriptome. Thus, positive fold-change values are genes transcribed at higher levels when the genus is highly transcriptionally active (compared to other metatranscriptomes). DESeq2 was also used to identify differentially transcribed genes for each

MAG between T0 and T24 samples in high tide (HT) dark incubations [23]. Since T24 samples were the “treatment” condition in DESeq2, positive fold-change values here are genes transcribed at higher levels in T24 compared to T0 samples. In all DESeq2 analyses, genes with Benjamini-Hochberg adjusted *p*<0.1 were counted as having significantly different transcription.

### QUANTITATIVE PCR

DNA samples from SAB field campaigns in 2014 and 2017 [24,54] were used as templates for qPCR reactions targeting the *Ca*. Poseidoniales 16S rRNA gene. Samples included the variety of shelf habitats (inshore, nearshore, mid-shelf, shelf-break, and oceanic as previously defined [24]; Fig. S1) and multiple depths when possible. DNA from the entire microbial community (no prefiltration) was extracted as previously described [54]. Primers were GII-554-f [55] and Eury806-r [56] with cycling conditions as previously reported [57] (Table S2). Reactions (25 μL, triplicate) used iTaq Universal Green SYBR Mix (Bio-Rad, Hercules, CA) in a C1000 Touch Thermal Cycler/CFX96 Real-Time System (Bio-Rad, Hercules, CA). Each plate included a no-template control and a standard curve (serial dilutions of a linearized plasmid containing a previously-sequenced, cloned amplicon). Abundance of *Ca*. Poseidoniales 16S rRNA genes was compared to published bacterial 16S rRNA gene abundance from the same samples [24,54]. Regional variability of gene abundance was assessed using a one-way ANOVA and a post-hoc HSD test as described above. Model II regressions of log-transformed qPCR data were estimated using the lmodel2 R package [58] as previously described [54]. All plots were constructed with anvi’o or the ggplot2 R package [59].

## RESULTS

### EURYARCHAEOTAL MAGs

SIMO Bins 19-2 and 31-1 were estimated as 82.5-92.3% and 77.5-96.2% complete, respectively, with redundancy <0.6% [25]. Phylogenomics placed both in the putative family *Ca*. Thalassarchaeaceae (MGIIb) and genera O1 (SIMO Bin 19-2) and O3 (SIMO Bin 31-1; Fig. S2). Phylogenomic groupings were generally consistent with previous findings [16,17].

Both SIMO MAGs contained a proteorhodopsin gene. Presence of a methionine residue at position 315 suggested absorption of green light [60], and both proteorhodopsin genes grouped in “Archaea Clade B” [11,17,61] (Fig. S3). Both MAGs included partial or complete pathways indicating aerobic heterotrophic growth, such as glycolysis, the TCA cycle, and electron transport chain components (Table S3). Pathways for metabolism of compounds such as fatty acids, peptides, and proteins were also present, as were transport systems and metabolic pathways for amino acids and nucleotides.

### DOMINANT GENERA IN FIELD METATRANSCRIPTOMES

There were significant seasonal differences in transcript recruitment by the combined set of *Ca*. Poseidoniales MAGs (*F*_3,28_=4.9, *p*=0.007. most summer samples had >10^10^ *Ca*. Poseidoniales transcripts L^−1^, significantly more than in other seasons (Fig. 1A,C; Table S4). The diversity of transcriptionally-active *Ca*. Poseidoniales also changed seasonally. Genera O1 and O3 accounted for most reads mapped from summer samples (typically 89.5-99.5% of *Ca*. Poseidoniales reads), with most mapping to O1 (Fig. 1A,B; Fig. S4; Table S4). HT (and not low tide; LT) metatranscriptomes from July 2014 also had a moderate fraction of reads (37.5-39.6%) mapped to *Ca*. Poseidoniaceae. In contrast to summer samples, November 2008 and May 2009 transcripts were predominantly O3, while those from February 2009 and October 2014 were mostly *Ca*. Poseidoniaceae genera M, L1, or L2 (Fig. 1A,B, Table S4).

### HIGHLY TRANSCRIBED *CA*. POSEIDONIALES GENES

Many highly transcribed *Ca*. Thalassarchaeaceae (MGIIb) genes were involved in translation, transcription, replication/repair, or post-translation protein modification (Fig. 2), as expected by their fundamental role in cellular processes. Genes encoding proteins involved in energy production or conservation (ATPases, a pyrophosphate-energized proton pump, and proteorhodopsin) were also highly transcribed. Notably, the *aapJ* and *livK* genes, encoding ligand-binding receptors for L-amino acid and branch-chain amino acid transporters, respectively, were among the most highly transcribed genes in both *Ca*. Thalassarchaeaceae MAGs (Fig. 2, Table S3).

**Fig. 2.**
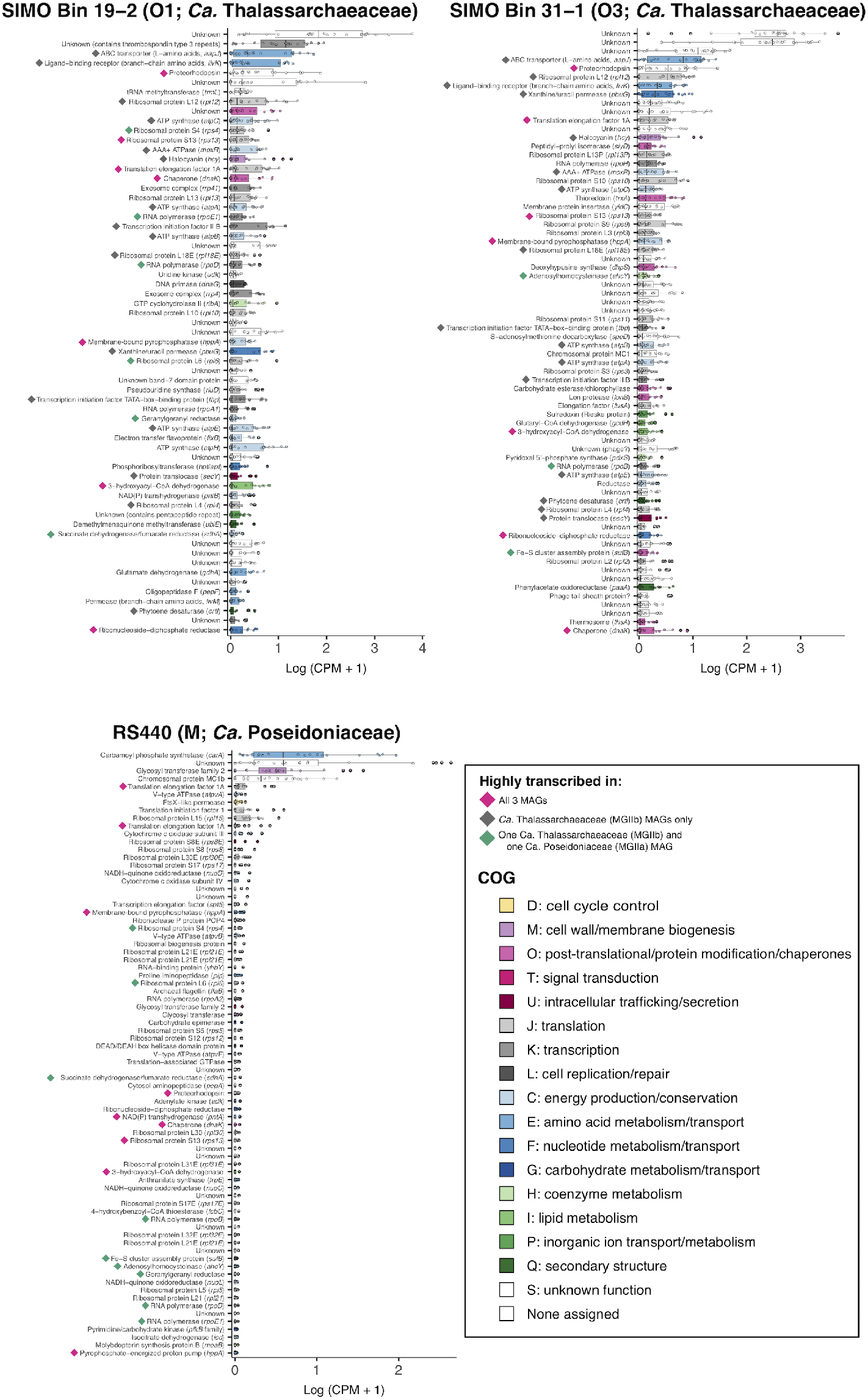
Boxplots of highly-transcribed MAG genes (top 5%) in Sapelo Island field metatranscriptomes. Overlain points show CPM for individual metatranscriptomes (*n*=24). Shading indicates COG functional category assigned by eggNOG-mapper (genes assigned to group S were similar to proteins of unknown function in the COG database, while genes with no COG assignment did not match proteins in the COG database). Diamonds indicate genes highly transcribed in all MAGs (pink), in *Ca*. Thalassarchaeaceae (MGIIb) MAGs only (gray), or in the *Ca*. Poseidoniaceae (MGIIa) MAG and one *Ca*. Thalassarchaeaceae MAG (green).

Many of the highly transcribed genes mapping to the *Ca*. Poseidoniaceae (MGIIa) MAG were not highly transcribed by *Ca*. Thalassarchaeaceae, including genes encoding a carbamoyl phosphate synthetase subunit (*carA*), a family 2 glycosyl transferase, chromosomal protein MC1b, and a ftsX-like permease. The *carA* gene had the highest median coverage of *Ca*. Poseidoniaceae genes across coastal metatranscriptomes (Fig. 2, Table S3). While both *Ca*. Thalassarchaeaceae MAGs also contained the carbamoyl phosphate synthetase genes, neither transcribed *carA* at high levels (Table S3).

Twelve genes were highly transcribed by *Ca*. Thalassarchaeaceae and not by *Ca*. Poseidoniaceae, including genes encoding ATP synthase, transcription initiation factor IIB, halocyanin, phytoene desaturase, protein translocase, xanthine/uracil permease, and receptors for amino acid transporters (Fig. 2, Table S3). Other than those encoding ribosomal proteins, only six genes were highly transcribed in all three MAGs: a chaperone protein, a ribonucleoside-diphosphate reductase, translation elongation factor 1A, 3-hydroxyacyl-CoA dehydrogenase, a membrane-bound pyrophosphatase (*hppA*), and proteorhodopsin (Fig. 2).

### DIFFERENTIAL GENE TRANSCRIPTION

We were interested in identifying genes with variable transcription levels when genera O1, O3, and M were highly transcriptionally active in the ocean. Twenty-three genes were differentially transcribed in O1-active samples (Fig. 3, Table S5), sixteen of which had higher abundance in O1-active metatranscriptomes compared to other metatranscriptomes. These highly transcribed genes encoded proteorhodopsin, two copper-containing redox proteins (halocyanin and plastocyanin), and proteins involved in lipid metabolism (3-hydroxyacyl-CoA dehydrogenase and oligosaccharyltransferase), nucleotide transport/metabolism (ribonucleotide-diphosphate reductase and xanthine/uracil permease), and amino acid transport (ligand-binding receptor for a L-amino acid transporter, *aapJ*). Differentially transcribed genes mapping to the O3 MAG were mostly depleted in O3-active metatranscriptomes and largely encoded proteins of unknown function; only the gene encoding ribosomal protein L12 was enriched in O3-active samples (Fig. S5, Table S5). Only four genes mapping to the M MAG were differentially transcribed in M-active metatranscriptomes. Annotated genes encoded chromosomal protein MC1b, an ATPase subunit, and a glycosyl transferase, which all had significantly fewer transcripts in M-rich samples (Fig. S6, Table S5). One gene of unknown function was enriched compared to other samples.

**Fig. 3.**
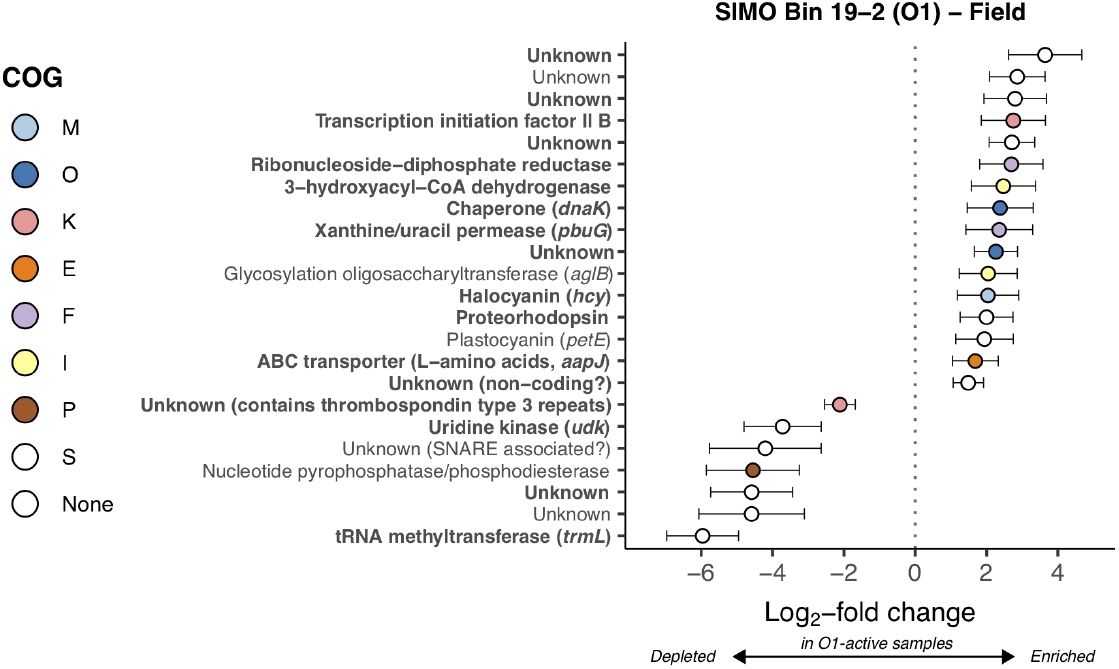
Log_2_-fold change of SIMO Bin 19-2 (genus O1) genes differentially transcribed in field metatranscriptomes where transcriptional activity of *Ca*. Poseidoniales was dominated by genus O1 (see Fig. 1, Table S4), calculated with DESeq2. Error bars show estimated standard error. Only genes with adjusted *p*-values <0.1 are shown. Color indicates COG functional category (see Fig. 2). Bold indicates genes in the top 5% median transcript coverage across field metatranscriptomes (Fig. 2).

### DARK INCUBATION METATRANSCRIPTOMES

Incubation had little effect on transcription by the dominant genera in LT samples (Fig. 4, Table S4). In contrast, there were distinct shifts in transcriptionally active populations during incubations of all HT samples. July HT metatranscriptomes initially contained 60.4-62.5% *Ca*. Thalassarchaeaceae (MGIIb) while hits from the corresponding T24 samples were 98.1-99.3% *Ca*. Thalassarchaeaceae, due to increased transcript hits to genus O1 (Fig. 4, Table S4). Likewise, October 2014 HT samples initially contained 65.0-66.6% hits to *Ca*. Poseidoniaceae (MGIIa) but changed to 78.3-98.8% hits to *Ca*. Thalassarchaeaceae at T24 due to an increase in hits to O1 (Fig. 4).

**Fig. 4.**
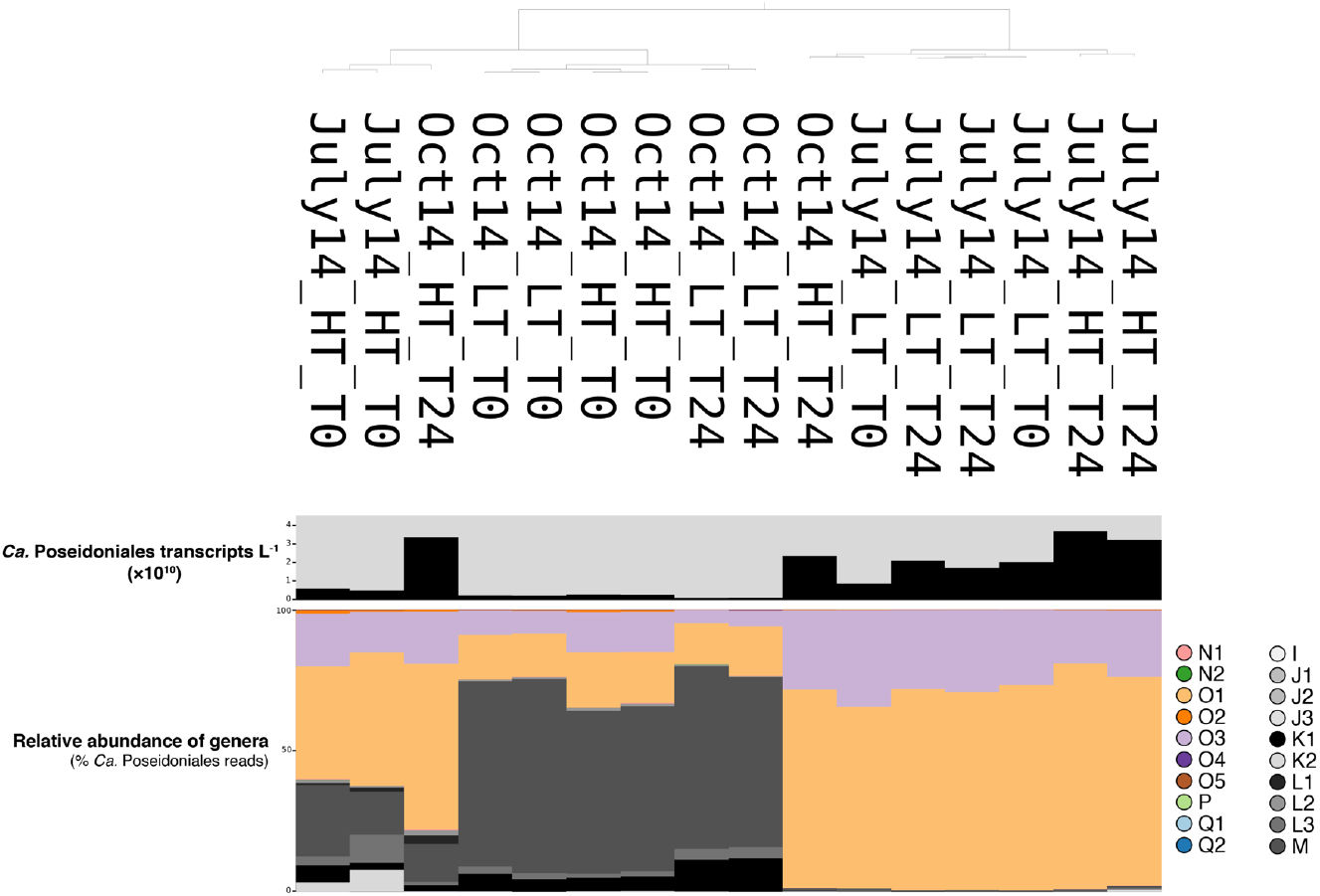
Comparisons of competitive read mapping to *Ca*. Poseidoniales genera at the beginning and end of 24-hour incubations of Sapelo Island water conducted by Vorobev et al. [23]. The dendrogram (top) shows grouping by similarity. The bar chart below the dendrogram is the abundance of *Ca*. Poseidoniales transcripts L^−1^. Stacked bar charts show the relative abundance of genera (% total *Ca*. Poseidoniales transcript reads), colored by genus.

DESeq2 identified 40 differentially transcribed genes mapping to the O1 MAG between HT T0 and T24 metatranscriptomes. Four O1 genes had higher transcription at T24, including xanthine/uracil permease (*pbuG*) and a ligand-binding receptor for a general amino acid transporter (*aapJ;* Fig. 5, Table S6). The 36 O1 genes transcribed at lower levels encoded proteins involved in repair of UV-damaged DNA, amino acid or nucleotide metabolism, coenzyme synthesis, peptidases or proteases, transcription, DNA replication, and lipid biosynthesis, as well as phytoene desaturase (*crtD*) and multiple subunits of pyruvate dehydrogenase (*pdhC, pdhA*). None of the genes mapping to the O3 or M MAGs were transcribed at significantly different levels between HT incubation timepoints (*p*>0.1 for all genes; Table S6).

**Fig. 5.**
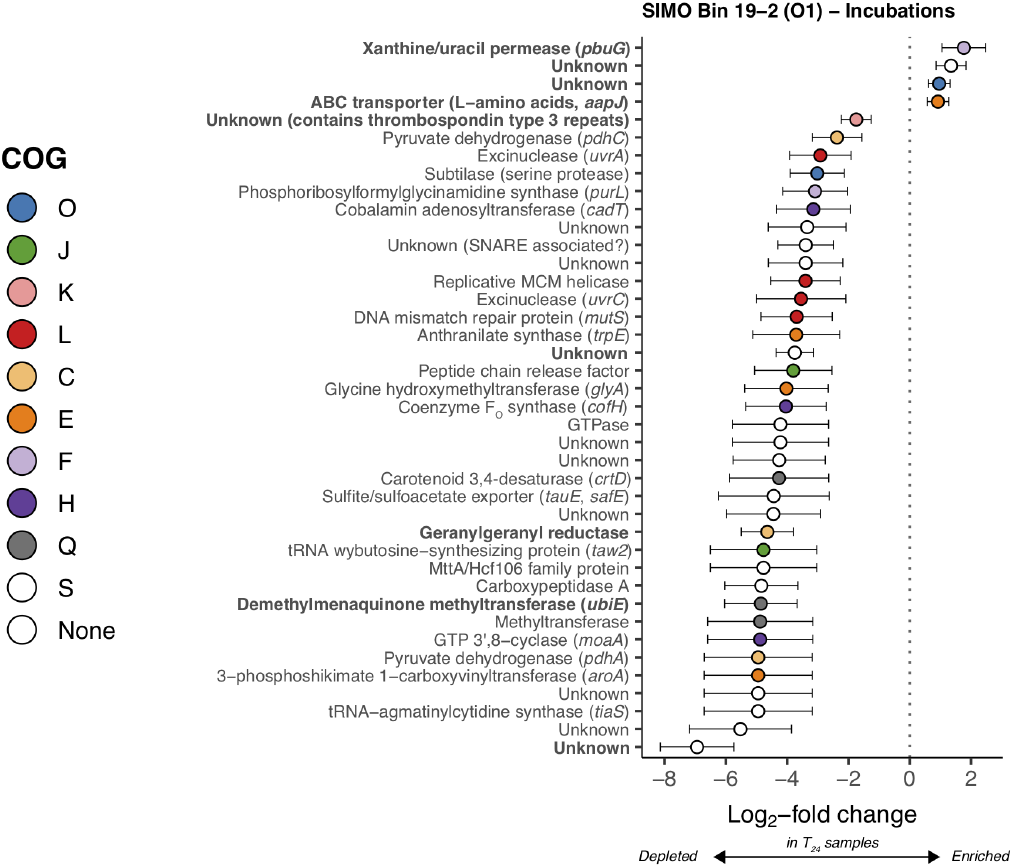
Log_2_-fold change of SIMO Bin 19-2 (genus O1) genes differentially abundant in T24 versus T0 metatranscriptomes from Sapelo Island high tide waters. Error bars show estimated standard error. Only genes with adjusted *p*-values<0.1 are shown. Color indicates COG functional category (see Fig. 2). Bold indicates genes in the top 5% median transcript coverage across field metatranscriptomes (Fig. 2).

### 16S rRNA GENE ABUNDANCE

*Ca*. Poseidoniales 16S rRNA genes were detected in all SAB DNA samples (*n*=208), with a range from 1.6×10^4^ to 7.6×10^8^ genes L^−1^ (Table S7). Standard curves for the *Ca*. Poseidoniales assay always had *r^2^*>0.99 (mean ± standard deviation: 0.99±0.001) and the mean efficiency was 93.4% (±2.0%; Table S2). When data from all cruises were combined, *Ca*. Poseidoniales genes were most abundant throughout inshore and nearshore waters and least abundant in shelf-break and oceanic samples (*F*_4,204_=18.5, *p*<0.001; Fig. 6A). There was a strong linear relationship between log-transformed bacterial and *Ca*. Poseidoniales 16S rRNA gene abundances, with bacterial abundances 2-3 orders of magnitude higher (Fig. 6B).

**Fig. 6.**
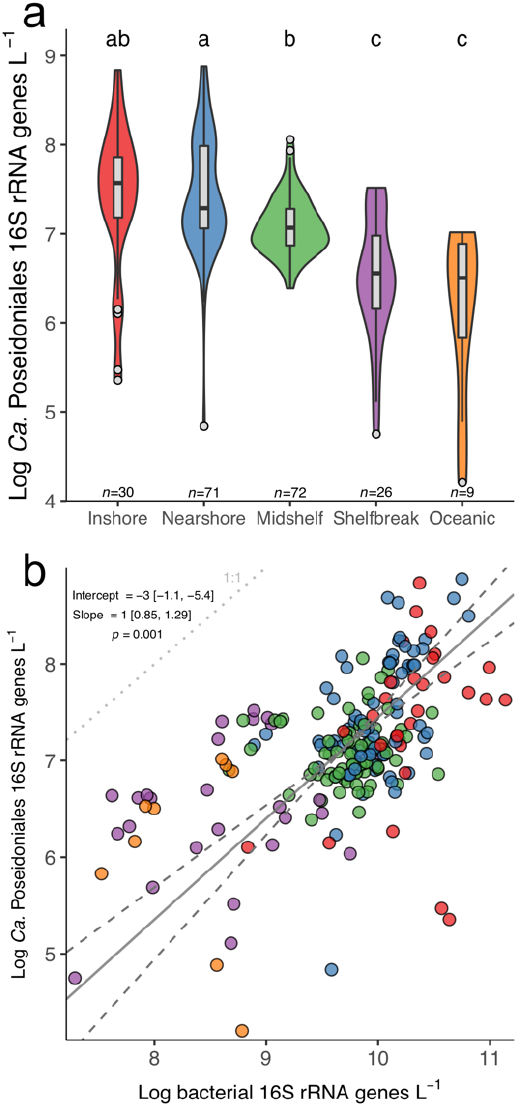
**A)** Violin plots of log-transformed *Ca*. Poseidoniales 16S rRNA gene abundances across regions in the SAB, with overlain boxplots. Width of the violin plot corresponds to data probability density. Color denotes sampling region. Letters above boxes denote post-hoc grouping according to Tukey’s HSD test. **B)** Scatterplot of bacterial and *Ca*. Poseidoniales 16S rRNA gene abundances in the SAB. The solid line shows the best fit of a model II (major axis) linear regression, with dashed lines showing a 95% confidence interval of the slope. Regression parameters are shown on the plot.

## DISCUSSION

### ABUNDANCE OF *CA*. POSEIDONIALES GENES IN THE SOUTH ATLANTIC BIGHT

Given typical *Ca*. Poseidoniales abundance of 10^6^-10^7^ genes or cells L^−1^ in oligotrophic waters [62–64] and 10^7^-10^8^ genes or cells L^−1^ in coastal waters [9,55,57,65–67], the abundance of their 16S rRNA genes in the coastal SAB (nearly 10^9^ genes L^−1^ in some samples) is among the highest measured in the ocean. Greater gene abundance in inshore, nearshore, and mid-shelf waters indicates *Ca*. Poseidoniales are more abundant over the shallow shelf than further offshore (Fig. 6A), matching clone library data from the SAB [24] and data from the Central California Current and the Black Sea [9,66]. The DOM in productive, turbid SAB coastal waters supports highly active heterotrophic microbial populations [68,69]. Our data suggest large populations of *Ca*. Poseidoniales are part of this heterotrophic community, and the correlation between *Ca*. Poseidoniales and bacterial abundance (Fig. 6B) suggests common factors influence the abundance of these two communities.

### DIVERSITY OF TRANSCRIPTIONALLY ACTIVE *CA*. POSEIDONIALES

While numerous studies have demonstrated high abundance of *Ca*. Poseidoniales in the coastal ocean (e.g., [57,67]), little is known about which clades are transcriptionally active in these regions. At Sapelo Island, the striking dominance of the *Ca*. Thalassarchaeaceae (MGIIb) genera O1 and O3 in summer samples, which also contained the highest levels of aggregate *Ca*. Poseidoniales transcripts (Fig. 1), indicates that *Ca*. Thalassarchaeaceae are most active during the summertime. Outliers to this pattern were July 2014 HT samples, which contained abundant *Ca*. Poseidoniaceae (MGIIa) transcripts (though *Ca*. Thalassarchaeaceae still comprised the majority of their *Ca*. Poseidoniales reads). A previous study found relatively high salinity and DOM enriched in marine-origin molecules over the mid-shelf SAB during July 2014 [70]. Our data suggest *Ca*. Poseidoniaceae were relatively active over the shelf during this time and were transported inshore during flood tides, leading to shifts in transcriptional diversity between LT waters (dominated by *Ca*. Thalassarchaeaceae) and HT waters (which included a higher number of *Ca*. Poseidoniaceae).

*Ca*. Poseidoniaceae (MGIIa) are the predominant euryarchaeal family in many coastal ecosystems, particularly in summer (e.g., [9,14,57,71,72]), but it is unclear what general patterns govern *Ca*. Poseidoniales distributions in coastal waters worldwide. Studies of *Ca*. Poseidoniales ecology often focus on distributions with depth, typically finding abundant *Ca*. Thalassarchaeaceae (MGIIb) in deeper waters and *Ca*. Poseidoniaceae (MGIIa) more prevalent in euphotic waters (e.g., [12,55,73–75]). A recent mapping of global ocean metagenome reads showed that coastal populations of *Ca*. Poseidoniales were primarily *Ca*. Poseidoniaceae, though *Ca*. Thalassarchaeaceae MAGs recruited a substantial number of reads from some coastal metagenomes [17]. Our data match this latter pattern, with *Ca*. Poseidoniales populations in surface waters off Sapelo Island dominated by highly active *Ca*. Thalassarchaeaceae (Fig. 1). The higher abundance of MAGs from genera O1 and O3 (also referred to as MGIIb. 12 and MGIIb.14) in some mesopelagic and coastal samples with relatively high temperature (~23–30°C; [17]) may explain the unusual pattern found in SAB waters: these genera peaked in summer at Sapelo Island, when water temperatures were 29–30°C (Table S4), suggesting they may be adapted to growth at relatively low light and high temperature.

### *CA*. POSEIDONIALES GENE TRANSCRIPTION PATTERNS

Sapelo Island metatranscriptome reads that mapped to *Ca*. Poseidoniales were analyzed in three ways:

1. We determined sets of “highly transcribed” genes mapping to MAGs of transcriptionally-active *Ca* Poseidoniales genera (5% of MAG genes with the highest median transcript coverage);
2. We identified genes mapping to *Ca* Poseidoniales MAGs that were differentially transcribed when its genus was highly active (≥50% of *Ca*. Poseidoniales transcripts in a sample; Fig. 1, Table S4);
3. We identified genes mapping to *Ca* Poseidoniales MAGs that were differentially transcribed at the beginning versus end of dark incubations. Since dark incubation separates indigenous microbes from light and sources of short-lived substrates, transcription of corresponding transporters and metabolic genes ceases during incubations as substrates are consumed. We therefore assume transcript depletion in T24 compared to T0 metatranscriptomes indicates genes that were transcriptionally active in the field [23]. This interpretation was bolstered by significant transcript depletions for genes involved in repairing UV damage to DNA (*uvrA*, *uvrC*, and *cofH*; Fig. 5), an expected result given alleviation of UV stress in dark incubations.

In the following sections, we synthesize these analyses to discuss *Ca*. Poseidoniales transcriptional activity related to DOM metabolism, transport/metabolism of amino acids and nucleotides, and basic energetic processes. Though lack of a cultivated representative limits the analysis to computationally-inferred functions, these data provide hypotheses regarding the activity of *Ca*. Poseidoniales families in the coastal ocean.

### PROTON GRADIENTS AND ELECTRON TRANSPORT

Our analysis revealed that *Ca* Poseidoniales genes involved in establishing transmembrane proton gradients were highly transcribed in our samples. Genes encoding proteorhodopsin were among the most highly transcribed by both *Ca*. Thalassarchaeaceae MAGs and were highly transcribed in O1-active samples (Figs. 2,3). Proteorhodopsins consist of a retinal chromophore linked to a transmembrane protein and use light energy to pump protons across the cell membrane [76]. The resulting energy can be coupled to ATP production or other chemiosmotic processes and often supports photoheterotrophy, though its function varies widely [61]. Proteorhodopsin genes are highly transcribed in the photic zone of both open ocean and coastal waters (e.g., [77–79]) and our data indicate coastal *Ca*. Poseidoniales conform to this pattern, consistent with recent evidence from other regions [21,80,81]. High transcription of proteorhodopsin supports the photoheterotrophic lifestyle hypothesized for *Ca*. Poseidoniales (e.g., [9,11,12,16,17]).

Since O1 proteorhodopsin transcript abundance did not differ between the beginning and end of dark incubations (Fig. 5), light may not regulate *Ca*. Thalassarchaeaceae proteorhodopsin transcription. However, depletion of O1 *crtD* (carotenoid 3,4-desaturase) transcripts during dark incubation (Fig. 5) suggests light may regulate retinal synthesis. Whether proteorhodopsin transcription responds to light varies among marine bacteria (e.g., [82,83]), and the function of constitutive transcription is not straightforward: while some bacteria use proteorhodopsin to produce ATP when carbon-limited [84], high amounts of proteorhodopsin in other bacteria can physically stabilize membranes even when inactive [85]. The high proteorhodopsin transcription in our data emphasizes, but provides little mechanistic clarification of, the physiological role for proteorhodopsin in *Ca*. Poseidoniales (Table 1).

**Table 1.**
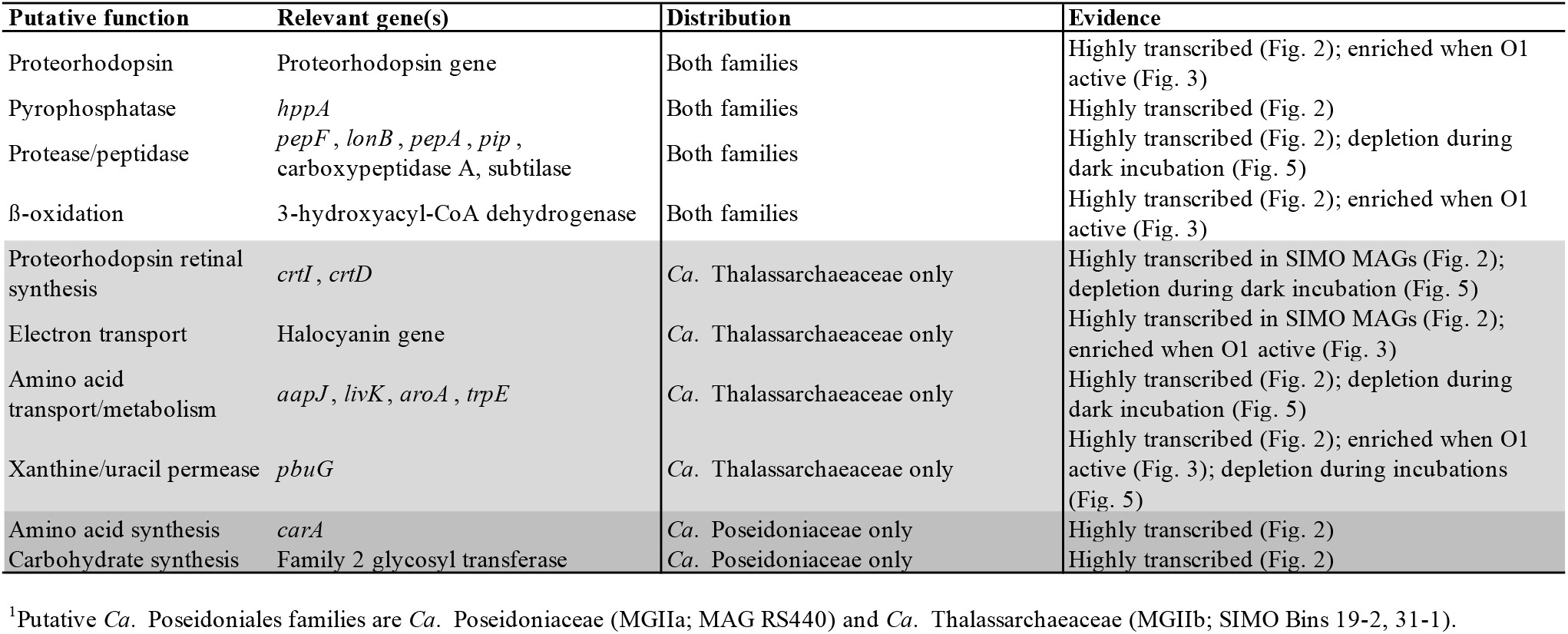
Transcriptional traits shared or distinct among *Ca*. Poseidoniales families.^1^

Numerous genes encoding putative electron transport proteins were highly transcribed by at least one *Ca*. Poseidoniales family (Fig. 2). Like proteorhodopsin, a halocyanin gene was among the most highly transcribed *Ca*. Thalassarchaeaceae genes and was enriched when genus O1 was transcriptionally active (Fig. 3). Halocyanins are involved in the electron transport chain and have been posited to increase the energy yield of aerobic respiration in *Ca*. Thalassarchaeaceae to stimulate rapid growth [12]. The similar transcription patterns of proteorhodopsin and halocyanin suggests proteorhodopsin activity in coastal *Ca*. Thalassarchaeaceae may function to increase growth rates during respiration, aiding rapid population growth when conditions permit.

The *hppA* gene from all *Ca* Poseidoniales MAGs was highly transcribed (Fig. 2; Table 1). This gene (which is found across many domains of life [86], including a diverse array of marine microbes [87]) putatively encodes a membrane-bound pyrophosphatase, which generates a proton gradient via hydrolysis of pyrophosphate, a byproduct of numerous cellular processes [86]. In metatranscriptomes from a phytoplankton bloom, enrichment of *hppA* transcripts suggested high pyrophosphate-based energy conservation in oligotrophic waters [87], and *hppA* transcripts were similarly abundant at night in an oligotrophic lake [88]. Although widespread in MAGs from *Ca*. Poseidoniales [16], *hppA* has not been previously recognized as a potentially important part of their metabolism. Our data suggest *Ca*. Poseidoniales may be capable of using pyrophosphatase (along with proteorhodopsin) to generate a protonmotive force (Table 1).

### POTENTIAL IMPORTANCE OF MARINE DOM IN *CA*. POSEIDONIALES METABOLISM

Although the T0 metatranscriptomes from summer versus fall were dominated by transcripts from different *Ca*. Poseidoniales families, a 24-hour dark incubation consistently favored transcription by *Ca*. Thalassarchaeaceae when samples were collected at HT (Fig. 4). This tidal stage-linked increase in *Ca*. Thalassarchaeaceae transcription could relate to differences in DOM availability between HT and LT, consistent with numerous studies implicating DOM in shaping *Ca*. Poseidoniales populations [9,10,89]. The HT Sapelo Island DOM pool is primarily of marine origin while LT DOM is more riverine- and marsh-derived [70], which may select for growth of different *Ca*. Poseidoniales families in the water masses present at different tidal stages. Furthermore, the depletion of transcripts encoding two pyruvate dehydrogenase subunits (*pdhA* and *pdhC*) during incubations (Fig. 5) suggests *Ca*. Poseidoniales were metabolizing phytoplankton photosynthate *in situ*. Alternatively, these tidal stage-driven transcriptional patterns may relate to differential light adaptation in populations originating in offshore versus nearshore waters. Inshore populations, potentially adapted to life in turbid waters, may increase transcription upon dark enclosure, whereas offshore populations (transported shoreward during flood tide) may be adapted to clearer waters and reduce transcription in dark conditions.

Multiple lines of evidence indicate coastal *Ca*. Poseidoniales may have been metabolizing proteins and fatty acids. High transcription of genes encoding proteases or peptidases from all *Ca*. Poseidoniales MAGs (Fig. 2) suggests likely metabolism of proteins or peptides by both families (Table 1). Furthermore, decreased transcription of protease genes mapping to the O1 MAG during dark incubations (Fig. 5) suggests protein metabolism by *Ca*. Thalassarchaeaceae may have been active *in situ* prior to incubation. While some of these genes could be involved in intracellular recycling (particularly lon protease and cytosol aminopeptidase), active protein metabolism is consistent with previous experiments demonstrating protein assimilation [10] and high transcription of *Ca*. Poseidoniales peptidase genes in other marine regions [15,21]. In addition to genes encoding protein catabolism, high transcription of the 3-hydroxyacyl-CoA dehydrogenase gene from all three MAGs (Fig. 2), and its enrichment in O1-active field samples (Fig. 3), suggests a likely importance of fatty acid metabolism for both *Ca*. Poseidoniales families (Table 1), matching widespread β-oxidation genes in *Ca*. Poseidoniales MAGs [16,17] and transcriptional data from the deep ocean [15].

### DISTINCT PATTERNS OF AMINO ACID AND NUCLEOTIDE UPTAKE AND METABOLISM

Transcription of *livK* and *aapJ* appears to differentiate *Ca*. Poseidoniales families in the coastal ocean (Table 1). These genes putatively encode ligand-binding receptors for ABC transporters: *aapJ* for a general L-amino acid transporter and *livK* for a branched-chain amino acid transporter [90]. Both are commonly present in *Ca*. Thalassarchaeaceae (MGIIb) but not *Ca*. Poseidoniaceae (MGIIa) [17] and were among the most highly transcribed *Ca*. Thalassarchaeaceae genes (Fig. 2). Previous studies noted high transcription of euryarchaeal *livK* and *aapJ* genes in the water column of the Red Sea [20,21], at the Mid-Cayman Rise [15], and throughout the Atlantic Ocean [22]. Our data suggest this activity was probably associated with *Ca*. Thalassarchaeaceae.

The *aapJ* and *livK* genes were collocated with genes putatively encoding the full transporters in the *Ca*. Thalassarchaeaceae MAGs (Table S3). Unfortunately, it is difficult to guess their substrates from sequence data alone: AAP transporters are typically capable of transporting a range of L-amino acids [90] while LIV transporters can be highly specific for leucine, specific for branched-chain amino acids, or transport diverse amino acids [90–92]. In soil bacteria grown under inorganic nitrogen limitation, elevated transcription of *aapJ* is linked to organic nitrogen use [93], but it is unclear whether this mechanism translates to *Ca*. Thalassarchaeaceae since amino acids could be used for numerous anabolic or catabolic processes. In addition to these binding proteins, the depletion of transcripts from O1 genes putatively involved in the shikimate pathway of aromatic amino acid synthesis (3-phosphoshikimate 1-carboxyvinyltransferase and anthranilate synthase) during incubations (Fig. 5) suggests *Ca*. Thalassarchaeaceae in this genus may have been synthesizing aromatic amino acids *in situ*.

The combination of high *pbuG* transcription by *Ca*. Thalassarchaeaceae (MGIIb) with the high numbers of O1 *pbuG* transcripts in O1-active samples and dark incubation endpoints (Figs. 2,3,5) suggests an important role for xanthine/uracil permease (the putative product of *pbuG*) in *Ca*. Thalassarchaeaceae metabolism. In some phytoplankton, *pbuG* is transcribed during nitrogen-stressed growth [94–96], potentially allowing access to DON. However, *pbuG* and xanthine dehydrogenase (*xdh*) are also transcribed when xanthine is catabolized by marine bacteria [97]. Both *Ca*. Thalassarchaeaceae MAGs contain putative xanthine dehydrogenase genes (*xdhC* and *yagS*; Table S3), suggesting the ability to catabolize xanthine (Table 1).

Transcription levels of *carA*, putatively encoding part of carbamoyl phosphate synthetase, appears to be a distinct trait of *Ca*. Poseidoniaceae (MGIIa): while all three MAGs contained this gene (Table S3), only *Ca*. Poseidoniaceae *carA* transcription was high. Since carbamoyl phosphate synthetase is a key enzyme for arginine and pyrimidine synthesis from bicarbonate [98], high *carA* transcription suggests these pathways may be important for *Ca*. Poseidoniaceae growth or survival, though it is not clear whether this transcription is related to synthesis of amino acids, nucleotides, or both.

## CONCLUSIONS

Our metatranscriptomic data and associated experiments provide a novel window into the activity of *Ca*. Poseidoniales families (formerly “MGIIa” and “MGIIb”). They indicate an important role for *Ca*. Thalassarchaeaceae (MGIIb) as coastal photoheterotrophs, particularly in warm waters. High transcription of proteorhodopsin and membrane-bound pyrophosphatase genes suggested common methods for establishing proton gradients. Furthermore, high transcription of genes involved in protein/peptide metabolism and β-oxidation of fatty acids confirmed peptide and lipid metabolism as a common trait. However, high transcription of *Ca*. Thalassarchaeaceae genes encoding amino acid binding proteins and nucleotide transporters suggests uptake of these substrates may distinguish the two families. These data confirm the importance of DOM metabolism by *Ca*. Poseidoniales and suggest a potential role for organic nitrogen in *Ca*. Thalassarchaeaceae metabolism.

## ACKNOWLEDGMENTS

We thank Qian Liu, Bradley Tolar, the Georgia Coastal Ecosystems LTER field team, the University of Georgia Marine Institute at Sapelo Island staff, and the captain and crew of the R/V *Savannah* for assistance with original field sampling. This work was funded in part by NSF OCE grant 1538677 and OPP grant 1643466 (to JTH), NSF Grant IOS1656311 (to MAM), NSF OCE grant 1832178 (to the Georgia Coastal Ecosystems LTER), as well as by resources and expertise from the Georgia Advanced Computing Resource Center, a partnership between the University of Georgia’s Office of the Vice President for Research and Office of the Vice President for Information Technology.

## CONFLICT OF INTEREST

The authors declare no competing financial interests.

## Notes

### Competing Interest Statement

The authors have declared no competing interest.

