## Supplemental text and figures for "Transcriptional activity differentiates families of Marine Group II *Euryarchaeota* in the coastal ocean"

1 **SUPPLEMENTARY FIGURES**
2 **Fig. S1**

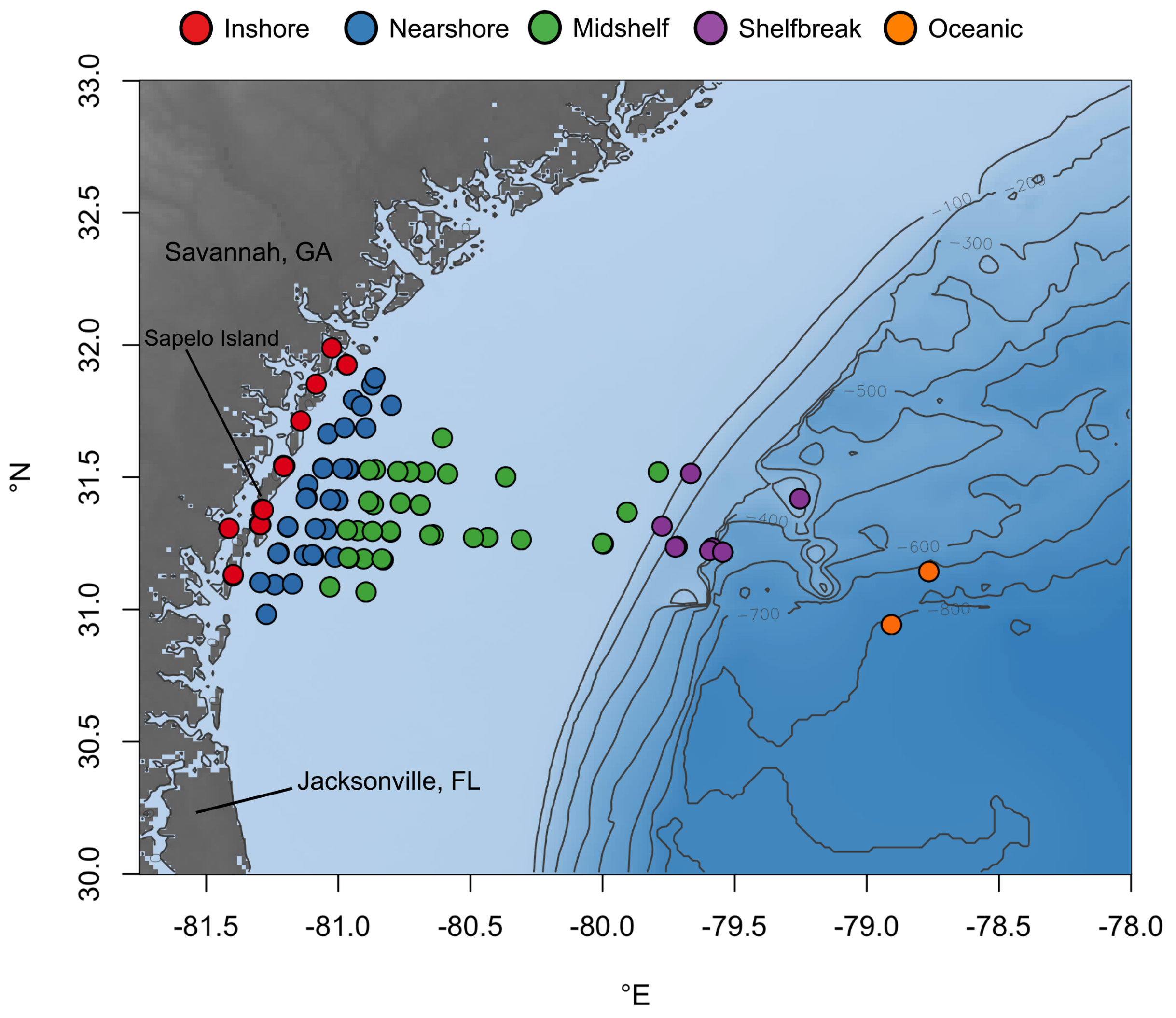


3

4 **Fig. S2**

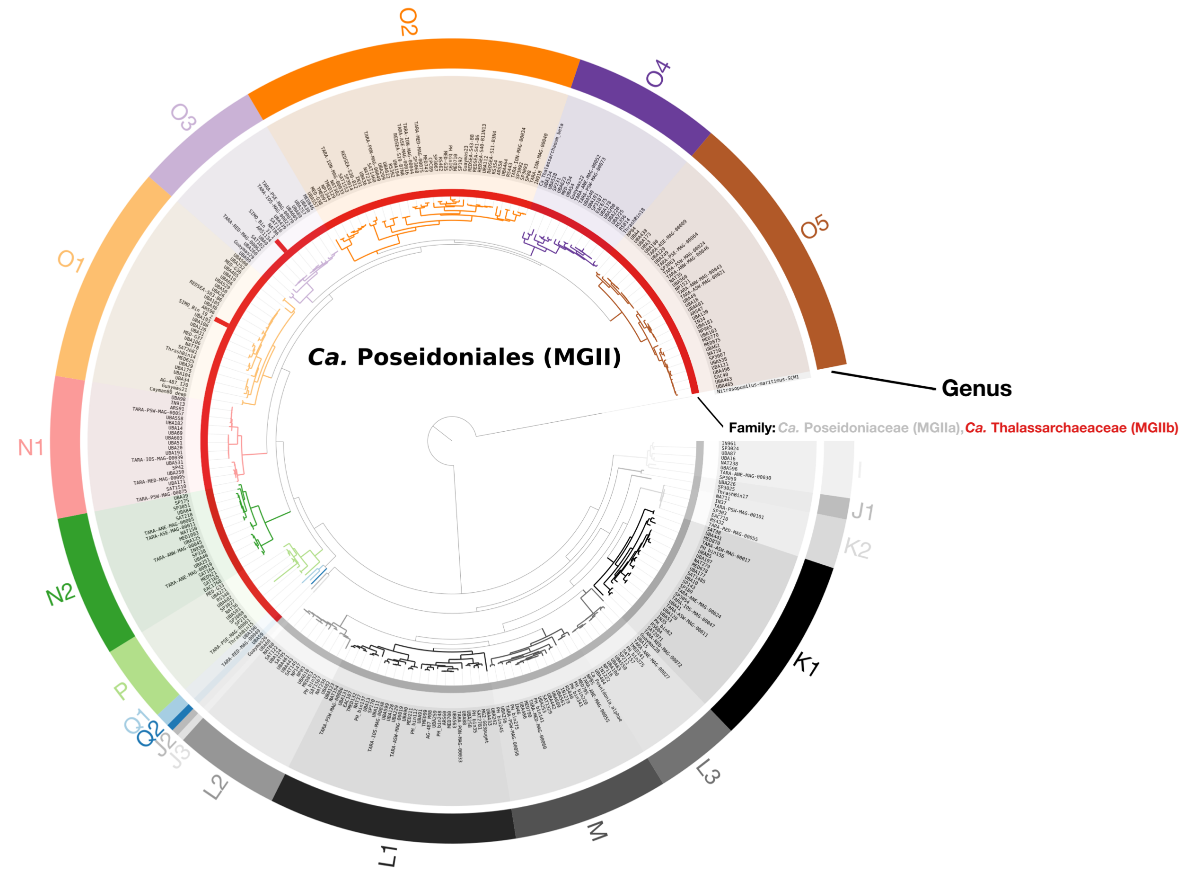
O2
O4
O3
O5
O1
Ca. Poseidoniales (MGII)
Genus
Family: Ca. Poseidoniaceae (MGIIa), Ca. Thalassarchaeaceae (MGIIb)
N1
N2
P
Q1
Q2
J2
J3
L2
L1
M
L3
K1
K2
J1
I

5

### Fig. S3

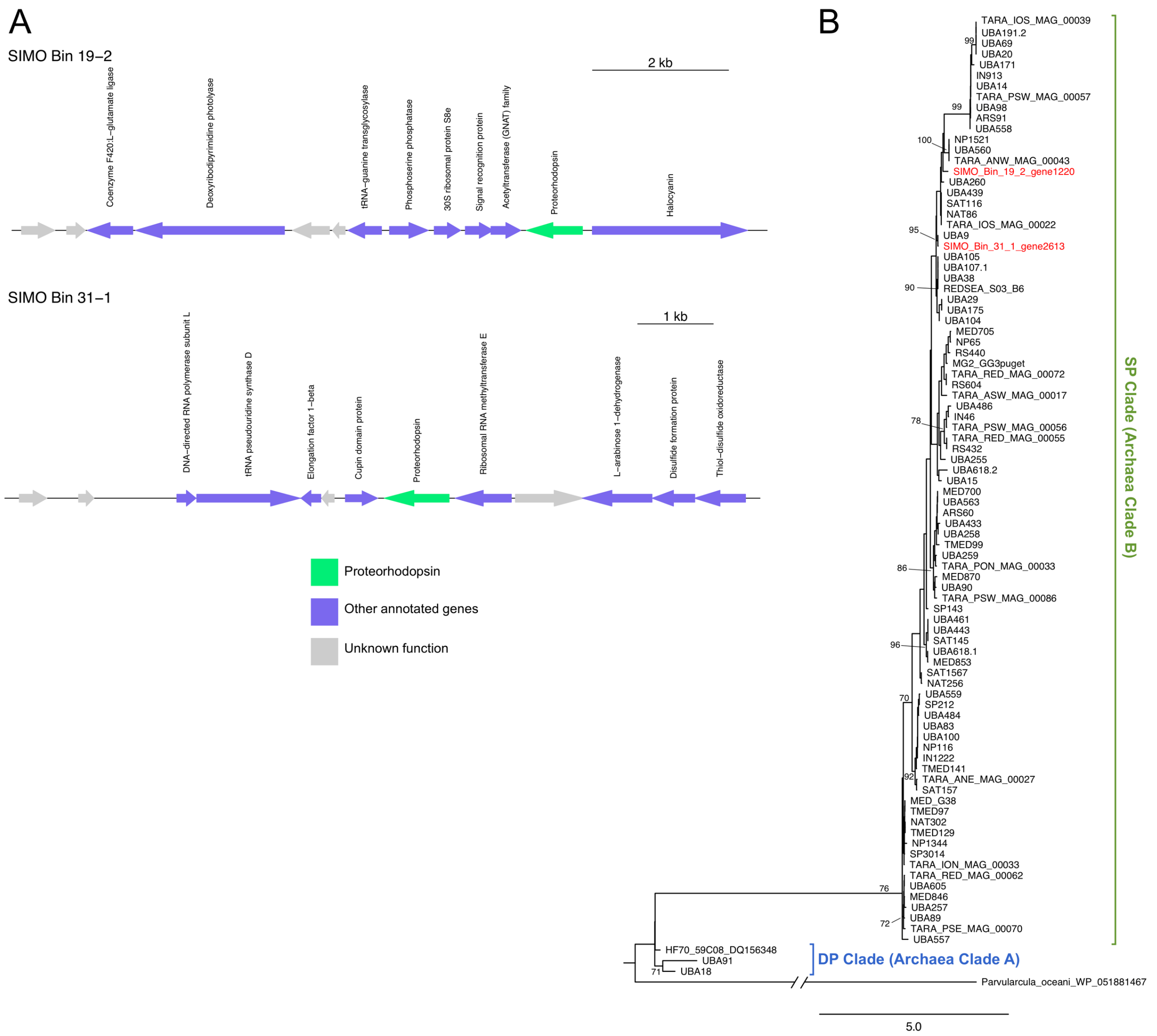
A
SIMO Bin 19–2
2 kb
Coenzyme F420:L–glutamate ligase
Deoxyribodipyrimidine photolyase
tRNA–guanine transglycosylase
Phosphoserine phosphatase
30S ribosomal protein S8e
Signal recognition protein
Acetyltransferase (GNAT) family
Proteorhodopsin
Halocyanin
SIMO Bin 31–1
1 kb
DNA–directed RNA polymerase subunit L
tRNA pseudouridine synthase D
Elongation factor 1–beta
Cupin domain protein
Proteorhodopsin
Ribosomal RNA methyltransferase E
L–arabinose 1–dehydrogenase
Disulfide formation protein
Thiol–disulfide oxidoreductase
Proteorhodopsin
Other annotated genes
Unknown function
B
TARA_IOS_MAG_00039
UBA191.2
UBA69
UBA20
UBA171
IN913
UBA14
TARA_PSW_MAG_00057
UBA98
ARS91
UBA558
NP1521
UBA560
TARA_ANW_MAG_00043
SIMO_Bin_19_2_gene1220
UBA260
UBA439
SAT116
NAT86
TARA_IOS_MAG_00022
UBA9
SIMO_Bin_31_1_gene2613
UBA105
UBA107.1
UBA38
REDSEA_S03_B6
UBA29
UBA175
UBA104
MED705
NP65
RS440
MG2_GG3puget
TARA_RED_MAG_00072
RS604
TARA_ASW_MAG_00017
UBA486
IN46
TARA_PSW_MAG_00056
TARA_RED_MAG_00055
RS432
UBA255
UBA618.2
UBA15
MED700
UBA563
ARS60
UBA433
UBA258
TMED99
UBA259
TARA_PON_MAG_00033
MED870
UBA90
TARA_PSW_MAG_00086
SP143
UBA461
UBA443
SAT145
UBA618.1
MED853
SAT1567
NAT256
UBA559
SP212
UBA484
UBA83
UBA100
NP116
IN1222
TMED141
TARA_ANE_MAG_00027
SAT157
MED_G38
TMED97
NAT302
TMED129
NP1344
SP3014
TARA_ION_MAG_00033
TARA_RED_MAG_00062
UBA605
MED846
UBA257
UBA89
TARA_PSE_MAG_00070
UBA557
HF70_59C08_DQ156348
UBA91
UBA18
Parvularcula_oceani_WP_051881467
99
99
100
95
90
78
86
96
70
92
76
72
71
SP Clade (Archaea Clade B)
DP Clade (Archaea Clade A)
5.0

**Fig. S4**

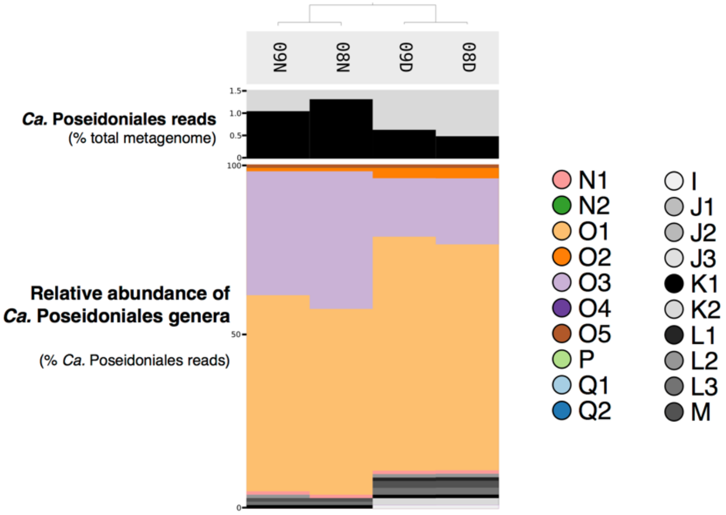
N60
N80
D60
D80
Ca. Poseidoniales reads
(% total metagenome)
1.5
1.0
0.5
0
100
Relative abundance of
Ca. Poseidoniales genera
50
(% Ca. Poseidoniales reads)
0
N1
N2
O1
O2
O3
O4
O5
P
Q1
Q2
I
J1
J2
J3
K1
K2
L1
L2
L3
M

**Fig. S5**

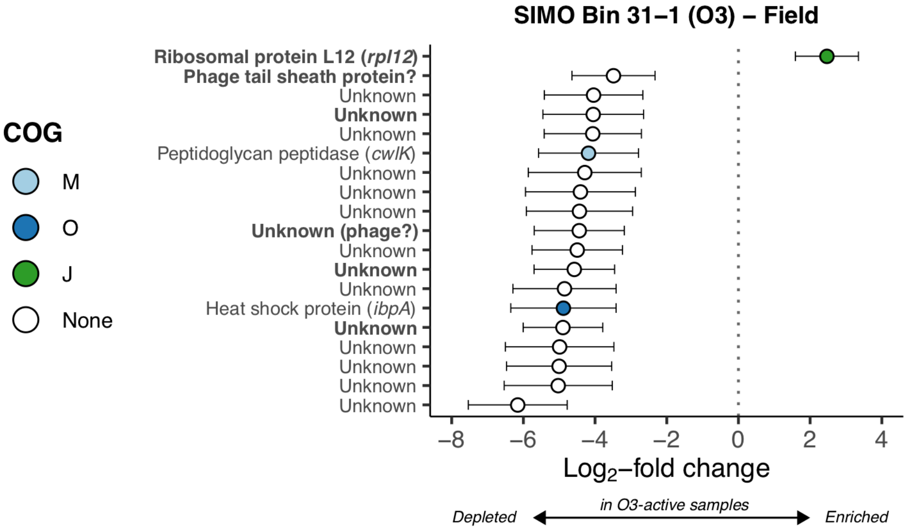
SIMO Bin 31–1 (O3) – Field
COG
M
O
J
None
Ribosomal protein L12 (rpl12)
Phage tail sheath protein?
Unknown
Unknown
Unknown
Peptidoglycan peptidase (cwlK)
Unknown
Unknown
Unknown
Unknown (phage?)
Unknown
Unknown
Unknown
Heat shock protein (ibpA)
Unknown
Unknown
Unknown
Unknown
Unknown
–8
–6
–4
–2
0
2
4
Log2–fold change
in O3-active samples
Depleted
Enriched

**Fig. S6**

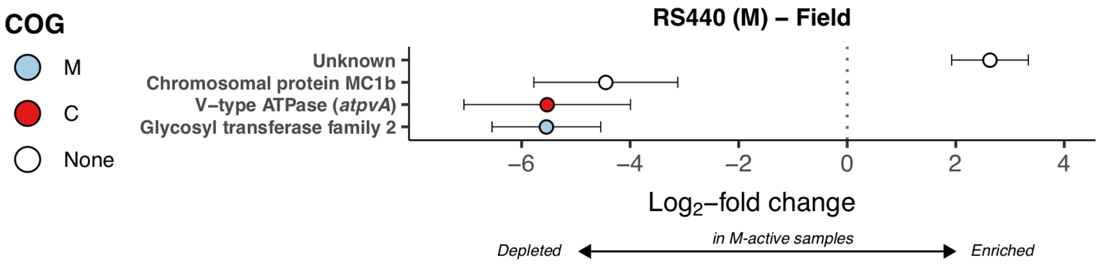
COG
M
C
None
RS440 (M) – Field
Unknown
Chromosomal protein MC1b
V–type ATPase (atpvA)
Glycosyl transferase family 2
−6
−4
−2
0
2
4
Log2–fold change
Depleted
in M-active samples
Enriched

### SUPPLEMENTARY FIGURE LEGENDS

**Fig. S1** Locations of SAB seawater collected for *Ca.* Poseidoniales qPCR analysis, constructed with the marmap R package [1]. For details of sampling locations see [2,3]. Sapelo Island (the location of metagenome and metatranscriptome collection) is indicated. Color indicates sampling region.

**Fig. S2** Phylogenomic tree of *Ca.* Poseidoniales MAGs, based on up to 16 concatenated ribosomal proteins. Red lines indicate the two SIMO MAGs. The outer circle shows genera based on [4]. *Nitrosopumilus maritimus* SCM1 is used as an outgroup.

**Fig. S3 A)** Maps of contigs containing proteorhodopsin genes (green) for SIMO Bins 19-2 (top) and 31-1 (bottom). Other annotated genes are blue and hypothetical proteins are gray. Contigs were illustrating using the genoPlotR R package [5]. **B)** Maximum likelihood amino acid phylogeny of proteorhodopsin genes. Genes from SIMO bins are shown in red. SP (Archaeal Clade B) and DP (Archaea Clade A) are marked in green and blue, respectively, adjacent to the tree. Branch labels show bootstrap support (100 ML replicates) of major clades with values >70; for clarity, support values for most inner clades are not shown. amino acid alignments were constructed using MUSCLE within Geneious [6] using a gap open penalty of -5. The evolutionary model was estimated with ProtTest3 [7] and was used to build a maximum likelihood tree using PhyML [8].

**Fig. S4** Relative abundance of genera in Sapelo Island metagenomes from summer 2008 and summer 2009 (*n*=4) [9]. The dendrogram (top) shows grouping by similarity. The bar chart shows the abundance of *Ca.* Poseidoniales transcripts $L^{-1}$ and the stacked bar charts show the relative abundance of genera (% total *Ca.* Poseidoniales transcripts), colored by genus. Since internal standards were not included in metagenomes, total *Ca.* Poseidoniales reads are shown as a percentage of the total metagenome.

**Fig. S5** $Log_2$-fold change of SIMO Bin 31-1 genes differentially transcribed in field metatranscriptomes where transcriptional activity of *Ca.* Poseidoniales was dominated by genus O3 (see Fig. 1), calculated with DESeq2. Error bars show estimated standard error. Only genes with adjusted *p*-values<0.1 are shown. Color indicates COG functional category (see Fig. 2). Bold indicates genes in the top 5% of median transcript coverage across field metatranscriptomes (Fig. 2).

**Fig. S6** $Log_2$-fold change of RS440 genes differentially transcribed in field metatranscriptomes where transcriptional activity of *Ca.* Poseidoniales was dominated by genus M (Fig. 1), calculated with DESeq2. Error bars show estimated standard error. Only genes with adjusted *p*-values<0.1 are shown. Color indicates COG functional category (see Fig. 2). Bold indicates genes in the top 5% of median transcript coverage across field metatranscriptomes (Fig. 2).

### SUPPLEMENTARY TABLE DESCRIPTIONS

**Table S1**

MAGs used in phylogenomics and competitive read mapping analyses. Clade assignments are from phylogenomics (Fig. S2).

**Table S2**

Information about primers, cycling conditions, and standard curve results from qPCR runs.

**Table S3**

Annotation and metatranscriptome coverages for genes in SIMO Bin 19-2, SIMO Bin 31-1, and RS440. Annotations using the MEROPS, dbCAN2, and TCDB databases used HMMER to search against database PFAM libraries; HMMER output is shown. Coverage (calculated with anvi'o) was normalized by dividing by the total number of reads in the metatranscriptome.

**Table S4**

Sample information for metatranscriptomes used in this study (see [10,11]) and the number, relative abundance, and absolute abundance of transcripts mapping to *Ca.* Poseidoniales MAGs.

**Table S5**

Differential transcription results from DESeq2 for field data. For each MAG, transcripts per gene were compared between samples where the respective genus was highly active versus samples where it was not (see Fig. 1). BH=Benjamini-Hochberg. Genes are numbered according to Table S3, with descriptions shown for genes with significantly differential transcription.

**Table S6**

Differential transcription results from DESeq2 for dark incubation data [11]. For each MAG, transcripts per gene were compared between $T_{24}$ and $T_0$ samples from high tide incubations (in which *Ca.* Poseidoniales transcript abundance changes; see Fig. 4). BH=Benjamini-Hochberg. Genes are numbered according to Table S3, with descriptions shown for genes with significantly differential transcription.

**Table S7**

Sampling information and gene quantities for SAB samples used in qPCR analysis. Locations, bacterial, and thaumarchaeal 16S rRNA quantities are reproduced from [2,3].
